## SupplementaryInformation for "Deep autoencoder for interpretable tissue-adaptive deconvolution and cell-type-specific gene analysis"

### Contents

**Supplementary Figure 1: Detailed comparison on pseudo-bulk datasets in the "normal" scenario.** 2
**Supplementary Figure 2: Detailed comparison on pseudo-bulk datasets in the "rare" scenario.** 2
**Supplementary Figure 3: Detailed comparison on three PBMC real datasets in the "similar" scenario.** 3
**Supplementary Table 1: Relations between defined cell types and existing cell types in original datasets.** 3
**Supplementary Figure 4: Deconvolution of immune cell subsets.** 4
**Supplementary Figure 5: Overall performance of all tested methods on five real datasets.** 4
**Supplementary Figure 6: Detailed comparison between Scaden and TAPE.** 5
**Supplementary Figure 7: Gene concordance of TAPE and CSx.** 5
**Supplementary Figure 8: Volcano plots of DEGs calculated from bulk GEPs and inferred GEPs.** 6
**Supplementary Figure 9: DEG detection would be affected by similar cell types.** 6
**Supplementary Figure 10: Comprehensive tests for TAPE and CIBERSORTx in four scenarios.** 7
**Supplementary Table 2: TAPE's performance is affected by variance cut-off.** 7
**Supplementary Figure 11: Data distribution after preprocessing.** 8
**Supplementary Table 3: Hyperparameters tuning for Scaden.** 8
**Supplementary Table 4: Hyperparameters tuning for RNAsieve.** 8
**Supplementary Table 5: Hyperparameters tuning for Music.** 9
**Supplementary Table 6: Hyperparameters tuning for DWLS.** 9
**Supplementary Table 7: Hyperparameters tuning for CIBERSORTx.** 9
**Supplementary Table 8: Performance summary of TAPE and SOTA methods.** 10
**Supplementary Table 9: Performance summary of TAPE and CIBESORTx on the DEG detection task.** 10

---

[*]Contribute equally

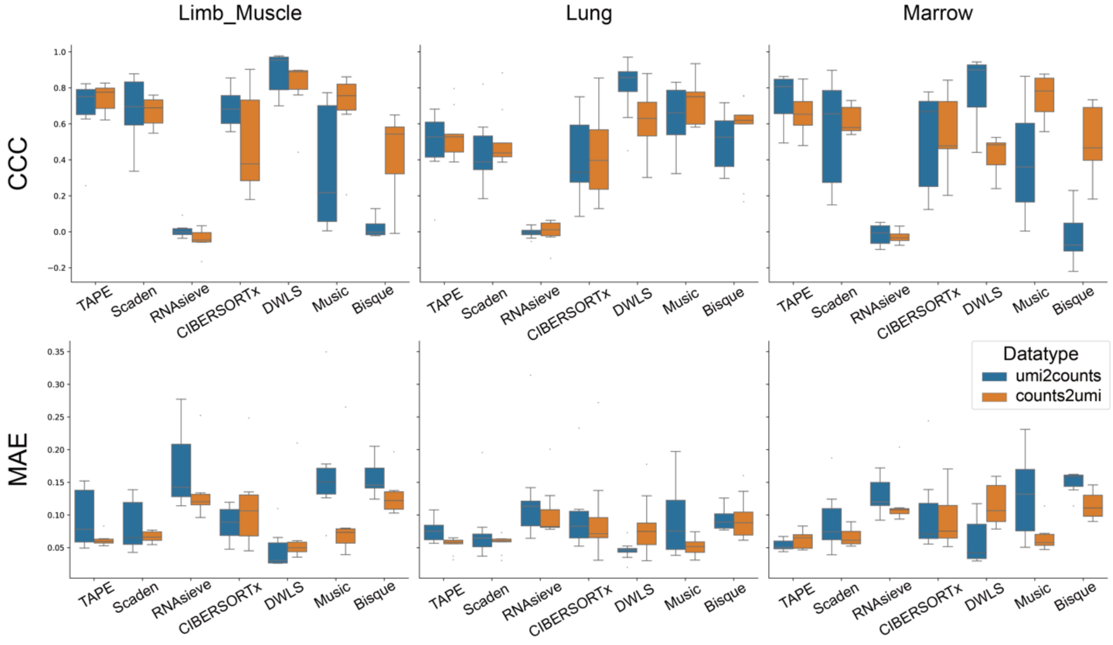


**Supplementary Figure 1: Detailed comparison on pseudo-bulk datasets in the "normal" scenario.** Each box contains performances of all the cell types in a certain tissue, sample sizes for each box in "Limb_Muscle", "Lung", and "Marrow" are 6, 9, and 7 respectively. In this figure, the boxes represent interquartile range (IQR) while the solid line represents the median. The whiskers extend to points that lie within 1.5 IQRs of the lower and upper quartile, and then observations that fall outside this range are displayed as points independently. Datatype refers to the cross-platform experiments. "umi2counts" means using single-cell profile from UMI-based data as reference to predict pseudo-bulk data constructed from counts-based single-cell profile and vice versa for "counts2umi". We can see many statistical methods suffers from batch effects if the datatype is exchanged. Source data are provided as a Source Data file.

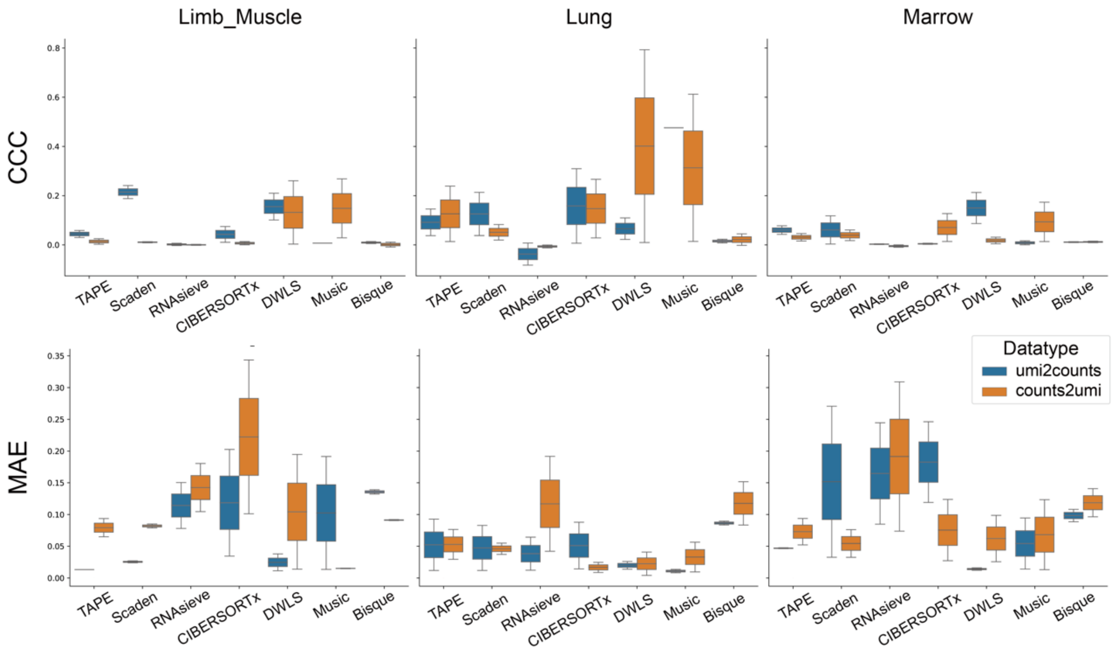


**Supplementary Figure 2: Detailed comparison on pseudo-bulk datasets in the "rare" scenario.** The figure settings are similar to Supplementary Figure 1 except that only rare cell types are considered in the three simulated datasets. Sample size for each box in each tissue consistently equals to 2. Similar to Supplementary Figure 1, the boxes represent IQR while the solid line represents the median. The whiskers extend to points that lie within 1.5 IQRs of the lower and upper quartile, and then observations that fall outside this range are displayed as points independently. All the methods can not achieve an appealing prediction power for rare cell types. Even though TAPE can achieve a relatively good performance on MAE (for example on umi2counts datatype, TAPE has the smallest average MAE on Limb_Muscle(0.013) and Marrow(0.047)), TAPE has worse performance on CCC in comparison with other methods, which indicates that TAPE needs further improvement in predicting a good correlation for rare cell types. Meanwhile, we have to point out that, for all the methods, a CCC value below 0.3 seems inadequate to show the deconvolution problem being resolved in the 'rare' scenario. Source data are provided as a Source Data file.

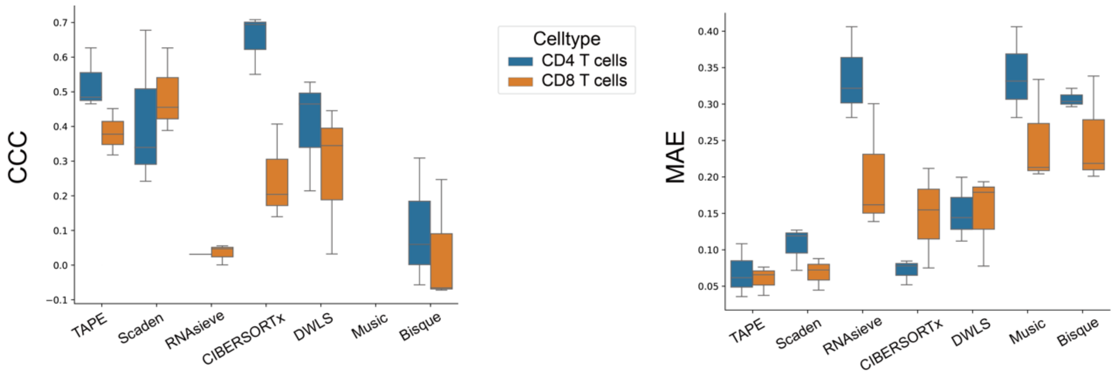


**Supplementary Figure 3: Detailed comparison on three PBMC real datasets in the "similar" scenario.** In all the three real PBMC datasets we considered, CD4 T cells and CD8 T cells exist in all of them. So we investigated the prediction performance for both of them across three PBMC datasets. Sample size for each box in the figure equals to 5. The boxes represent IQR while the solid line represents the median. The whiskers extend to points that lie within 1.5 IQRs of the lower and upper quartile, and then observations that fall outside this range are displayed as points independently. The results show that TAPE is the best algorithm for distinguishing similar cell types and has stable performance in the two cell types. Source data are provided as a Source Data file.

**Supplementary Table 1: Relations between defined cell types and existing cell types in original datasets.** Notice that we merge some cell types to make the categories identical.

| Defined cell types | data8k cell types | monaco's cell types |
|---|---|---|
| NK | CD16+ NK cells<br>NK cells | NK |
| Monocytes | Classical monocytes<br>Non-classical monocytes | Monocytes C<br>Monocytes N<br>Monocytes L |
| mDC | DC1<br>DC2 | mDCs |
| pDC | pDC | pDCs |
| Naïve B | Naïve B cells | B Naïve<br>B Exhausted |
| Memory B | Memory B cells | B SM<br>B NSM |
| MAIT | MAIT cells | MAIT |
| Naïve CD8 T | Tcm/Naive cytotoxic T cells | T CD8 Naive |
| Naïve CD4 T | Tcm/Naive helper T cells | T CD4 Naïve |
| non-Naïve CD4 T | Tem/Effector helper T cells | Tfh<br>Th1<br>Th1/Th17<br>Th17<br>Th2<br>T CD4 TE |
| non-Naïve CD8 T | Tem/Temra cytotoxic T cells<br>Tem/Trm cytotoxic T cells | T CD8 CM<br>T CD8 EM<br>T CD8 TE |
| Treg | Regulatory T cells | Tregs |
| Unknown | HSC/MPP | Progenitors<br>Plasmablasts<br>T gd Vd2<br>Tdg non-Vd2<br>Neutrophils LD<br>Basophiles LD |

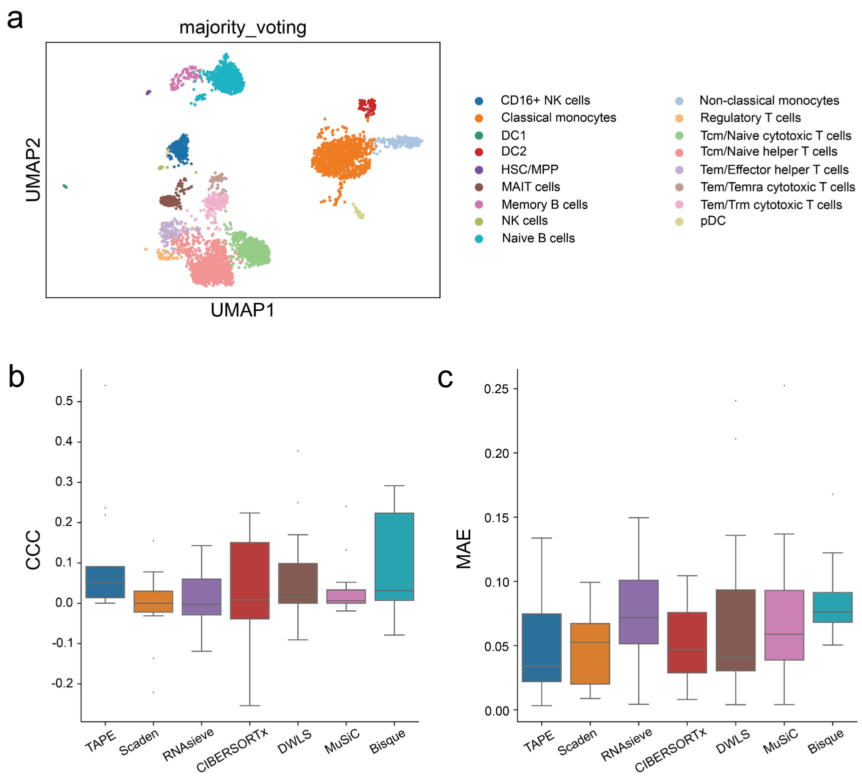


**Supplementary Figure 4: Deconvolution of immune cell subsets. a.** The annotated cell types of data8k datasets. This is produced from the pipeline of CellTypist [1]. Only cells with a confidence score greater than 0.8 were selected. **b,c.** Deconvolution performance of current methods on Monaco's dataset. Only TAPE predicts positive CCC for all cell types. Source data are provided as a Source Data file.

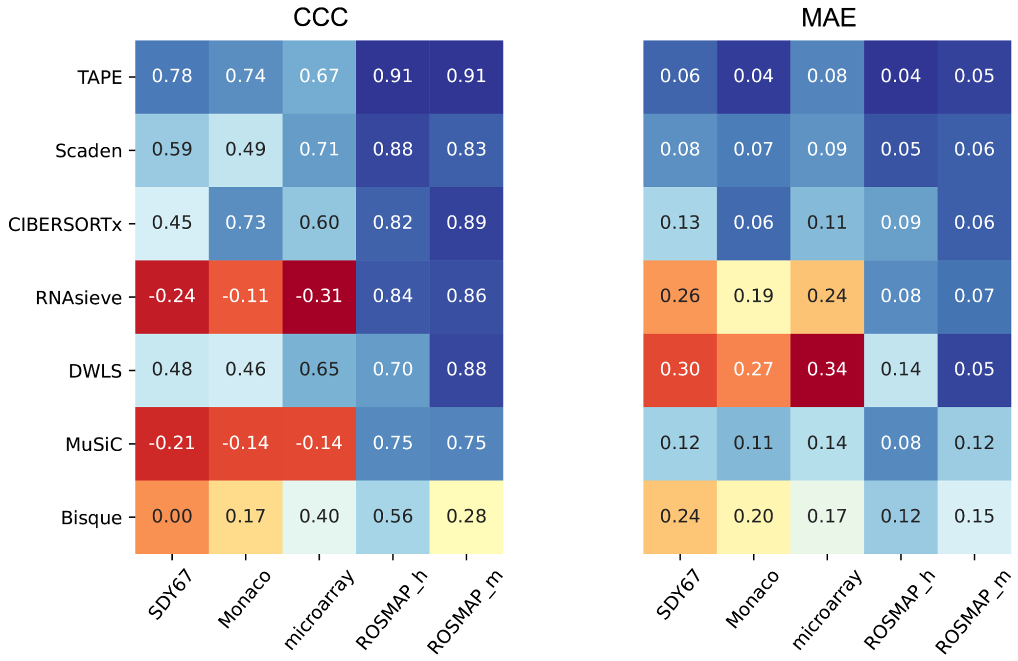


**Supplementary Figure 5: Overall performance of all tested methods on five real datasets.** The overall performance is calculated by all the data points of a dataset. Source data are provided as a Source Data file.

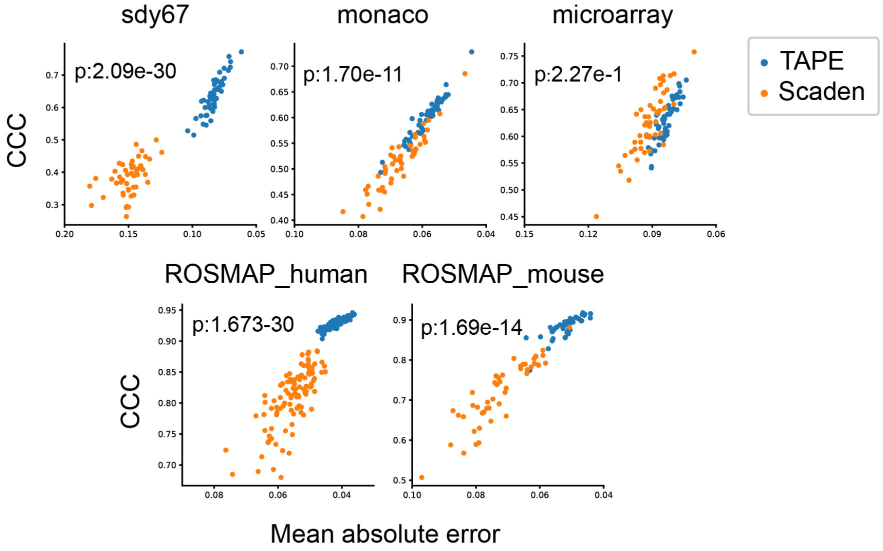


**Supplementary Figure 6: Detailed comparison between Scaden and TAPE.** A detailed comparison between the performance of Scaden and TAPE on 5 real bulk datasets using 50 different random seeds. The dots on the upper right represent better performance than the dots on the bottom left. Statistical significance is measured by a two-sided t-test which examines the distance from each of the two sets of points to the upper right corner. Source data are provided as a Source Data file.

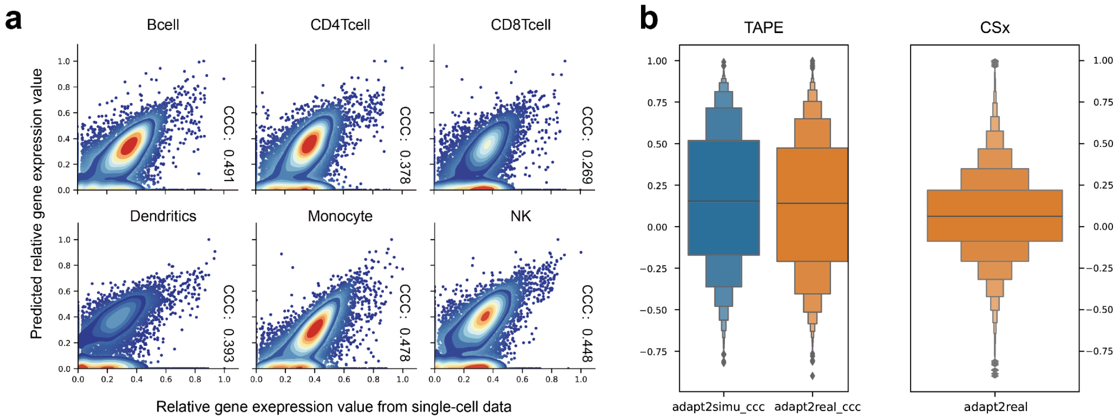


**Supplementary Figure 7: Gene concordance of TAPE and CSx.** a Concordance between the predicted relative gene expression value in real bulk data and the relative gene expression value in single-cell data of CSx. b Gene level CCC of TAPE and CSx (median CCC of TAPE and CSx in adapt2real scenario are 0.2171 and 0.0627, respectively). Source data are provided as a Source Data file.

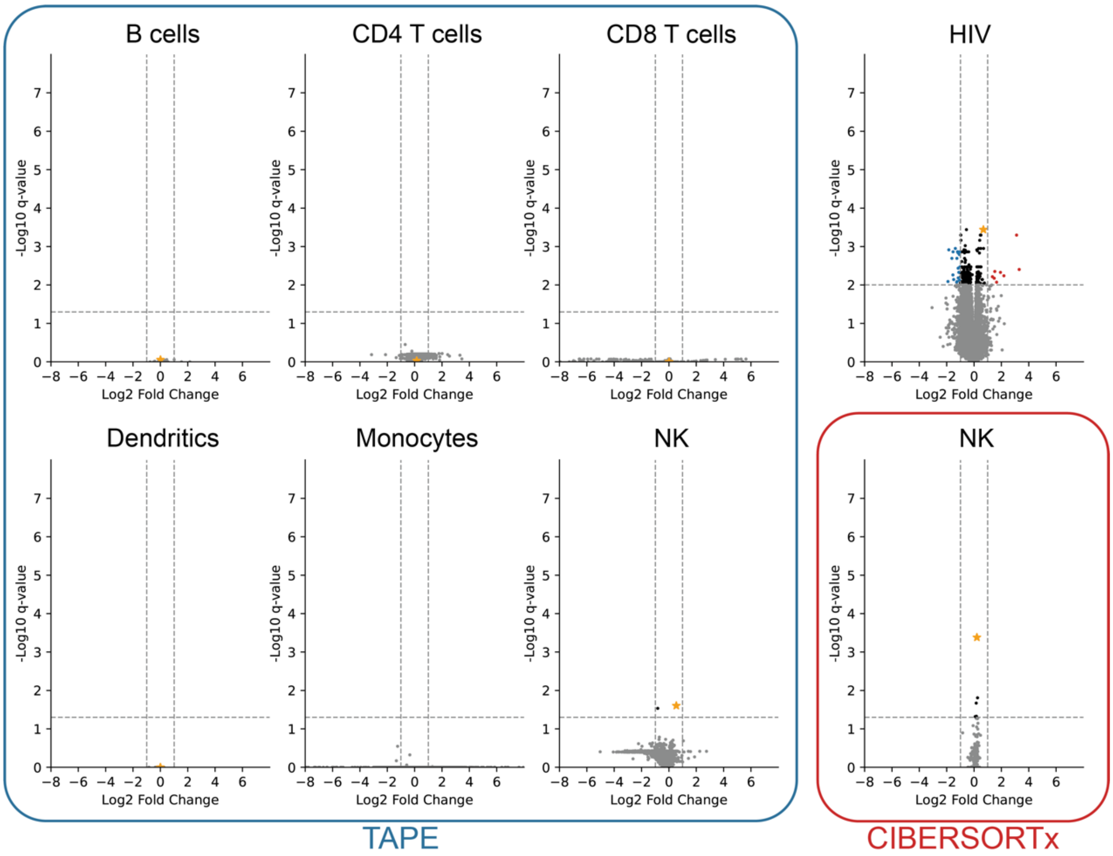


**Supplementary Figure 8: Volcano plots of DEGs calculated from bulk GEPs and inferred GEPs.** The q-value refers to the p-value adjusted by the false discovery rate. The orange stars refer to the RAB11FIP5 gene. The dash lines refers to the widely used p-value and foldchange criterions (log2(foldchange) > 1, q-value< 0.05 for inferred GEPs and q-value< 0.01 for real bulk GEPs). The DEGs of real bulk are detected using DESeq2 (without filtering out non-related conditions) [2], and the DEGs of inferred GEPs are detected by two-sided t-test. Both CIBERSORTx and TAPE can predict RAB11FIP5 as DEG in NK cells properly, but both methods can not infer a proper foldchange for DEG. Source data are provided as a Source Data file.

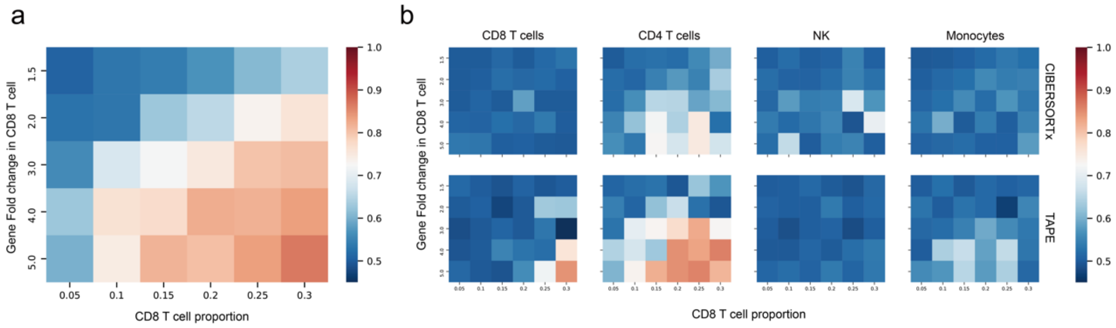


**Supplementary Figure 9: DEG detection would be affected by similar cell types (100 randomly selected DEGs with similar cell types). a.** Differentially expressed genes detected from simulated bulk RNA-seq data. The color indicates the AUROC value, red means better classification performance. Each row means different up-regulated foldchanges of randomly selected genes in CD8 T cells. Each column means CD8 T cell proportion in simulated bulk data. **b.** Differentially expressed genes detected by CIBERSORTx and TAPE in different cell types. DEGs should only be detected from CD8 T cells. But both methods prefer to think DEGs are from CD4 T cells. Source data are provided as a Source Data file.

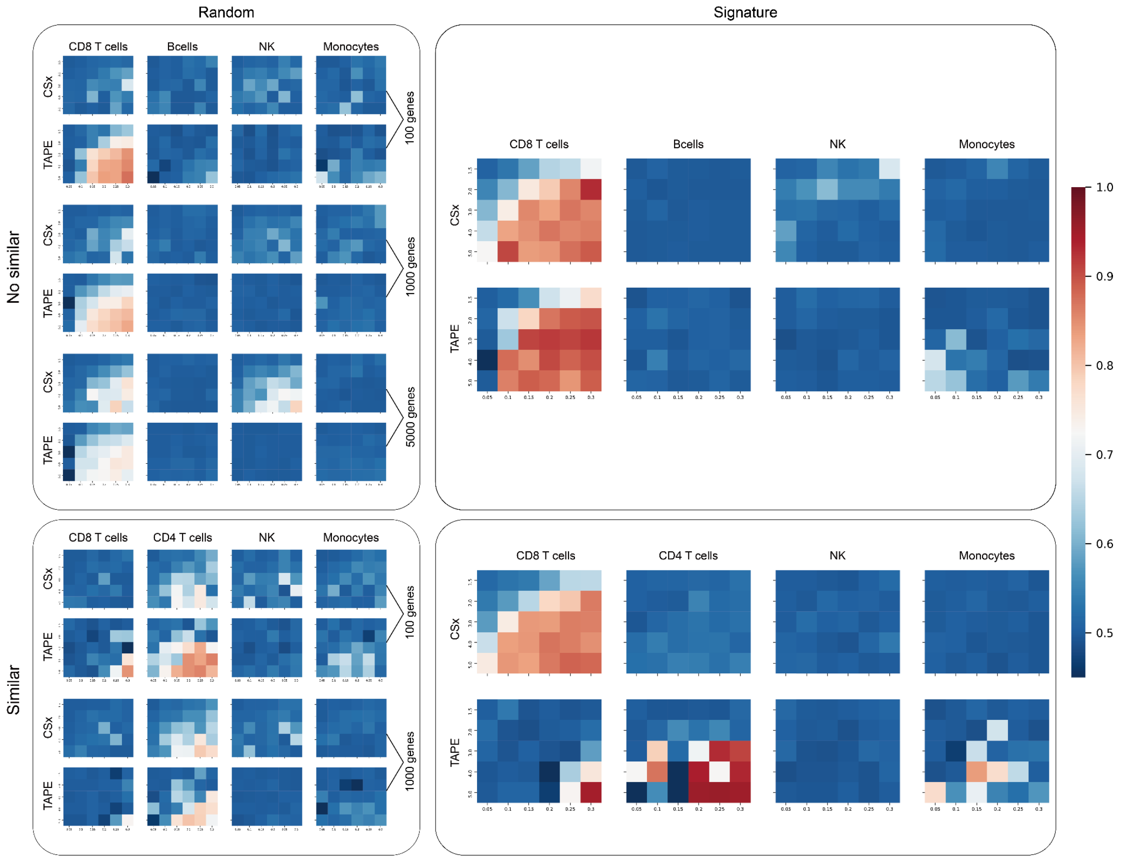


**Supplementary Figure 10: Comprehensive tests for TAPE and CIBERSORTx in four scenarios.** The upper left scenario uses randomly selected DEGs and it does not contain similar cell types in single-cell profiles. The number of DEGs ranges from 1,00 to 5,000. However, the number of DEGs is usually below 1,000 [3]. The second one is the “signature genes as DEGs without similar cell types” scenario which is located in the upper right. The bottom left area is the “randomly selected DEGs with similar cell types” scenario, and the bottom right one is the ”signature genes as DEGs with similar cell types” scenario. All the tests use AUROC as criteria, and the high AUROC value is expected to only appear in CD8 T cells. In the first scenario, TAPE is better than CIBERSORTx when the number of DEGs is below 5,000 (average AUROC in CD8 T cells for CSx and TAPE are 0.5578 and 0.6538 respectively). In the second scenario, both methods can achieve a good predictive power (average AUROC in CD8 T cells for CSx and TAPE are 0.7639 and 0.7611 respectively). In the third scenario, both methods can not distinguish DEGs from similar cell types well but TAPE’s performance is a little better (average AUROC in CD8 T cells for CSx and TAPE are 0.5146 and 0.5249 respectively). In the last scenario, CIBERSORTx behaves better than TAPE because of the incorporation of the signature matrix (average AUROC in CD8 T cell for CSx and TAPE are 0.7466 and 0.5336 respectively). Source data are provided as a Source Data file.

**Supplementary Table 2: TAPE’s performance is affected by variance cut-off.**

| **Fractions of genes left after cutoff** | 0.99 | 0.80 | 0.60 | 0.50 | 0.40 | 0.20 | 0.05 |
|---|---|---|---|---|---|---|---|
| **Number of genes left after cutoff** | 16599 | 16529 | 13200 | 10718 | 8076 | 3489 | 739 |
| **overall CCC** | 0.28 | 0.28 | 0.42 | 0.43 | 0.59 | 0.56 | 0.33 |
| **overall MAE** | 0.12 | 0.12 | 0.10 | 0.10 | 0.07 | 0.07 | 0.09 |

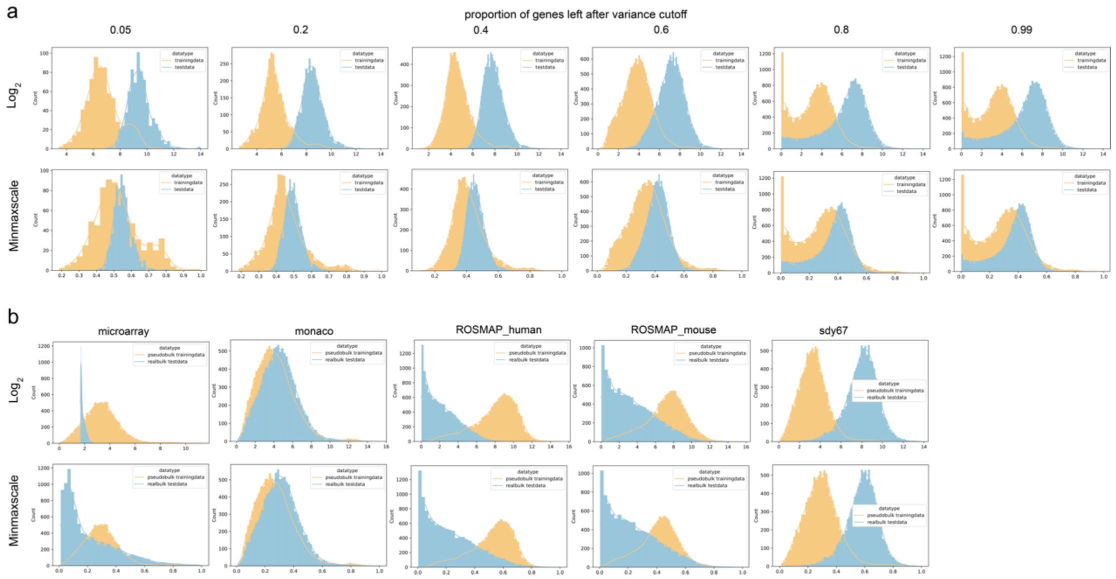


**Supplementary Figure 11: Data distribution after preprocessing.** In the data preprocessing step, our model needs to filter out low variance genes. a. Data distribution after filtering out different proportions of low variance genes. The test was conducted on the pseudo-bulk test: "Lung umi2counts". b. Data distribution of all the real bulk tests. Source data are provided as a Source Data file.

**Supplementary Table 3: Hyperparameters tuning for Scaden.**

| | | | | | | | | | |
|---|---|---|---|---|---|---|---|---|---|
| **parameters** | batch size | **128** | 64 | 64 | 64 | 64 | 128 | 128 | 128 |
| | learning rate | **1.00E-04** | 1.00E-04 | 1.00E-04 | 1.00E-05 | 1.00E-05 | 1.00E-04 | 1.00E-05 | 1.00E-05 |
| | steps | **5000** | 2000 | 5000 | 2000 | 5000 | 2000 | 2000 | 5000 |
| **metrics** | CCC_overall | 0.49 | 0.47 | 0.48 | 0.56 | 0.51 | 0.49 | 0.51 | 0.53 |
| | MAE_overall | 0.07 | 0.07 | 0.08 | 0.06 | 0.07 | 0.07 | 0.07 | 0.07 |
| | CCC_average | 0.37 | 0.32 | 0.32 | 0.33 | 0.31 | 0.31 | 0.33 | 0.31 |
| | MAE_average | 0.07 | 0.07 | 0.08 | 0.06 | 0.07 | 0.07 | 0.07 | 0.07 |

**Supplementary Table 4: Hyperparameters tuning for RNAsieve.**

| | | | | | | | | | | | | | |
|---|---|---|---|---|---|---|---|---|---|---|---|---|---|
| **parameters** | trim_percent | **0.02** | 0.10 | 0.05 | 0.01 | 0.02 | 0.10 | 0.05 | 0.01 | 0.02 | 0.10 | 0.05 | 0.01 |
| | gene_thresh | **0.20** | 0.20 | 0.20 | 0.20 | 0.10 | 0.10 | 0.10 | 0.10 | 0.30 | 0.30 | 0.30 | 0.30 |
| **metrics** | CCC_overall | -0.10 | -0.08 | -0.08 | -0.10 | -0.06 | -0.07 | -0.08 | -0.06 | -0.11 | -0.10 | -0.11 | -0.10 |
| | MAE_overall | 0.19 | 0.17 | 0.17 | 0.19 | 0.18 | 0.20 | 0.20 | 0.18 | 0.18 | 0.18 | 0.18 | 0.18 |
| | CCC_average | nan | nan | nan | nan | nan | nan | nan | nan | nan | nan | nan | nan |
| | MAE_average | 0.19 | 0.17 | 0.17 | 0.19 | 0.18 | 0.20 | 0.20 | 0.18 | 0.18 | 0.18 | 0.18 | 0.18 |

**Supplementary Table 5: Hyperparameters tuning for Music.**

| | | | | | | | | | | | | | | | | | | | | | |
|---|---|---|---|---|---|---|---|---|---|---|---|---|---|---|---|---|---|---|---|---|---|
| **Parameters** | nu | **0.0001** | 0.0000 | 0.0010 | 0.0100 | 0.1000 | 0.0001 | 0.0000 | 0.0010 | 0.0100 | 0.1000 | 0.0001 | 0.0010 | 0.0100 | 0.1000 | 0.0000 | 0.0001 | 0.0000 | 0.0010 | 0.0100 | 0.1000 |
| | centered | **FALSE** | FALSE | FALSE | FALSE | FALSE | TRUE | TRUE | TRUE | TRUE | TRUE | FALSE | FALSE | FALSE | FALSE | FALSE | TRUE | TRUE | TRUE | TRUE | TRUE |
| | normalized | **FALSE** | FALSE | FALSE | FALSE | FALSE | FALSE | FALSE | FALSE | FALSE | FALSE | TRUE | TRUE | TRUE | TRUE | TRUE | TRUE | TRUE | TRUE | TRUE | TRUE |
| **Metrics** | CCC_overall | -0.0275 | 0.0249 | -0.0262 | 0.1967 | 0.2373 | 0.0986 | 0.0821 | 0.1222 | 0.2324 | 0.2219 | -0.0165 | -0.0650 | -0.0924 | -0.0898 | 0.0044 | -0.0165 | 0.0280 | 0.0491 | 0.0302 | -0.0162 |
| | MAE_overall | 0.1796 | 0.1769 | 0.1683 | 0.1435 | 0.1731 | 0.1885 | 0.1954 | 0.1696 | 0.1496 | 0.1901 | 0.1739 | 0.1741 | 0.1771 | 0.1756 | 0.1712 | 0.1739 | 0.1821 | 0.1739 | 0.1786 | 0.1911 |
| | CCC_average | NaN | NaN | NaN | NaN | NaN | NaN | NaN | NaN | NaN | NaN | NaN | NaN | NaN | NaN | NaN | NaN | NaN | NaN | NaN | NaN |
| | MAE_average | 0.1796 | 0.1769 | 0.1683 | 0.1435 | 0.1731 | 0.1885 | 0.1954 | 0.1696 | 0.1496 | 0.1901 | 0.1739 | 0.1741 | 0.1771 | 0.1756 | 0.1712 | 0.1739 | 0.1821 | 0.1739 | 0.1786 | 0.1911 |

**Supplementary Table 6: Hyperparameters tuning for DWLS.**

| | | | | | |
|---|---|---|---|---|---|
| **parameters** | p-value cutoff | **0.0100** | 0.0100 | 0.0500 | 0.0500 |
| | diff cutoff | **0.5** | 0.5 | 0.5 | 0.5 |
| | flavor | **MAST** | Seurat | MAST | Seurat |
| **Metrics** | CCC_overall | 0.4056 | 0.4358 | 0.4056 | 0.1539 |
| | MAE_overall | 0.0979 | 0.1037 | 0.0979 | 0.2040 |
| | CCC_average | 0.2894 | 0.1515 | 0.2894 | 0.0554 |
| | MAE_average | 0.0979 | 0.1037 | 0.0979 | 0.2040 |

**Supplementary Table 7: Hyperparameters tuning for CIBERSORTx.**

| | | | | | | | | | | | |
|---|---|---|---|---|---|---|---|---|---|---|---|
| **parameters** | Kappa | **14.63** | 8.98 | 14.63 | 8.98 | 14.63 | 8.98 | 14.63 | 8.98 | 5 | 2 |
| | quantile normalization in generating signature matrix | **FALSE** | TRUE | FALSE | TRUE | FALSE | TRUE | FALSE | TRUE | FALSE | FALSE |
| | S-mode correction | **TRUE** | TRUE | TRUE | TRUE | FALSE | FALSE | FALSE | FALSE | TRUE | TRUE |
| | quantile normalization in deconvolution | **FALSE** | FALSE | TRUE | TRUE | FALSE | FALSE | TRUE | TRUE | FALSE | FALSE |
| **Metrics** | CCC_overall | 0.6167 | 0.4955 | 0.6108 | 0.5517 | 0.2260 | 0.0953 | 0.2389 | 0.1061 | 0.6817 | 0.5788 |
| | MAE_overall | 0.0683 | 0.0910 | 0.0659 | 0.0753 | 0.0866 | 0.0996 | 0.0875 | 0.0978 | 0.0603 | 0.0772 |
| | CCC_average | 0.3235 | 0.2254 | 0.3189 | 0.2542 | 0.3258 | 0.2504 | 0.3282 | 0.2594 | 0.3477 | 0.0772 |
| | MAE_average | 0.0683 | 0.0910 | 0.0659 | 0.0753 | 0.0866 | 0.0996 | 0.0875 | 0.0978 | 0.0603 | 0.0772 |

**Supplementary Table 8: Performance summary of TAPE and SOTA methods. Here we list the top3 methods in order in different scenarios and datasets.** The performance comparison between box plots is evaluated by two-sided t-test. The initial assumption is TAPE's performance is better than other methods. If $p > 0.5$, TAPE's performance is not better than other methods. The order is based on the p-value, a small p-value represents higher performance. In some scenarios with only two data points, we only compute the average value for comparison.

| Datatype | | Scenario | Dataset | Metrics | | | |
|---|---|---|---|---|---|---|---|
| | | | | CCC | | MAE | |
| | | | | Overall | P-value (used for comparing the box plot) | Overall | P-value (used for comparing the box plot) |
| Real-bulk | | | sdy67 | **TAPE**, Scaden, CSx | DWLS, Scaden, **TAPE** | **TAPE**, Scaden, MuSiC | **TAPE**, Scaden, CSx |
| | | | monaco | **TAPE**, CSx, Scaden | **TAPE**, CSx, Scaden | **TAPE**, CSx, Scaden | **TAPE**, CSx, Scaden |
| | | | microarray | Scaden, **TAPE**, DWLS | Scaden, CSx, **TAPE** | **TAPE**, Scaden, CSx | **TAPE**, Scaden, CSx |
| | | | rosmap_h | **TAPE**, Scaden, RNAsieve | **TAPE**, Scaden, MuSiC | **TAPE**, Scaden, MuSiC | **TAPE**, Scaden, MuSiC |
| | | | rosmap_m | **TAPE**, CSx, DWLS | RNAsieve, DWLS, **TAPE** | **TAPE**, DWLS, Scaden | **TAPE**, DWLS, Scaden |
| Pseudo-bulk | umi2counts | normal | Limb_Muscle | DWLS, CIBERSORTx, Scaden | DWLS, CIBERSORTx, Scaden | DWLS, Scaden, CIBERSORTx | Scaden, DWLS, CIBERSORTx |
| | | | Lung | DWLS, Bisque, **TAPE** | DWLS, Bisque, **TAPE** | DWLS, Scaden, **TAPE** | DWLS, Scaden, **TAPE** |
| | | | Marrow | DWLS, **TAPE**, Scaden | DWLS, **TAPE**, CIBERSORTx | **TAPE**, DWLS, Scaden | **TAPE**, DWLS, CIBERSORTx |
| | | rare* | Limb_Muscle | not applicable | Scaden, DWLS, **TAPE** | not applicable | **TAPE**, DWLS, Scaden |
| | | | Lung | not applicable | MuSiC, CIBERSORTx, Scaden | not applicable | MuSiC, DWLS, RNAsieve |
| | | | Marrow | not applicable | DWLS, Scaden, **TAPE** | not applicable | DWLS, **TAPE**, MuSiC |
| | | similar distinguishment* | Marrow | not applicable | **TAPE**, DWLS, CIBERSORTx | not applicable | **TAPE**, CIBERSORTx, Scaden |
| | | similar transferring* | Marrow | not applicable | DWLS, **TAPE**, Scaden | not applicable | DWLS, **TAPE**, Scaden |
| | counts2umi | normal | Limb_Muscle | **TAPE**, DWLS, Scaden | DWLS, **TAPE**, MuSiC | **TAPE**, Scaden, DWLS | **TAPE**, DWLS, Scaden |
| | | | Lung | MuSiC, **TAPE**, Scaden | MuSiC, DWLS, Bisque | MuSiC, **TAPE**, Scaden | MuSiC, **TAPE**, Scaden |
| | | | Marrow | MuSiC, **TAPE**, Scaden | MuSiC, **TAPE**, Scaden | **TAPE**, Scaden, MuSiC | **TAPE**, MuSiC, Scaden |
| | | rare* | Limb_Muscle | not applicable | MuSiC, DWLS, **TAPE** | not applicable | MuSiC, **TAPE**, Scaden |
| | | | Lung | not applicable | DWLS, MuSiC, CIBERSORTx | not applicable | CIBERSORTx, DWLS, MuSiC |
| | | | Marrow | not applicable | MuSiC, CIBERSORTx, Scaden | not applicable | Scaden, DWLS, MuSiC |
| | | similar distinguishment* | Marrow | not applicable | MuSiC, **TAPE**, CIBERSORTx | not applicable | **TAPE**, MuSiC, CIBERSORTx |
| | | similar transferring* | Marrow | not applicable | MuSiC, **TAPE**, Scaden | not applicable | MuSiC, **TAPE**, Scaden |

*only compute the average performance of all data points because of the small number of data points

**Supplementary Table 9: Performance summary of TAPE and CIBESORTx on the DEG detection task.** The performance is evaluated by the average AUROC in CD8 T cells. Since DEGs are only associated with different conditions which are not related to cell types' signature genes, we usually care about the case that DEGs are randomly selected.

| Scenario | DEG type | Similar cell type | DEG number | Performance |
|---|---|---|---|---|
| 1 | random | no | 100 | TAPE > CIBERSORTx |
| | | | 1000 | TAPE > CIBERSORTx |
| | | | 5000 | TAPE ~ CIBERSORTx |
| 2 | signature | no | about 200 | TAPE ~ CIBERSORTx |
| 3 | random | yes | 100 | both failed, TAPE > CIBERSORTx |
| | | | 1000 | both failed, TAPE > CIBERSORTx |
| 4 | signature | yes | about 150 | CIBERSORTx > TAPE |
